# Kupffer Cell Release of Platelet Activating Factor Drives Dose Limiting Toxicities of Nucleic Acid Nanocarriers

**DOI:** 10.1101/2020.02.11.944504

**Authors:** Meredith A. Jackson, Shrusti S. Patel, Fang Yu, Matthew A. Cottam, Evan B. Glass, Bryan R. Dollinger, Ella N. Hoogenboezem, Prarthana Patil, Danielle D. Liu, Isom B. Kelly, Sean K. Bedingfield, Allyson R. King, Rachel E. Miles, Alyssa M. Hasty, Todd D. Giorgio, Craig L. Duvall

## Abstract

*In vivo* nanocarrier-associated toxicity is a significant and poorly understood hurdle to clinical translation of siRNA nanomedicines. In this work, we demonstrate that platelet activating factor (PAF), an inflammatory lipid mediator, plays a key role in nanocarrier-associated toxicities, and that prophylactic inhibition of the PAF receptor (PAFR) completely prevents these toxicities. High-dose intravenous injection of siRNA-polymer nano-complexes (si-NPs) elicited acute, shock-like symptoms (vasodilation and vascular leak) in mice and caused a three-fold increase in blood PAF levels. PAFR inhibition completely prevented these toxicities, indicating PAF activity is a primary driver of systemic si-NP toxicity. Pre-treatment with clodronate liposomes fully abrogated si-NP-associated increases in blood PAF and consequent toxicities, suggesting that nanoparticle uptake by Kupffer macrophages is the source of PAF. Assessment of varied si-NP chemistries further confirmed that toxicity level correlated to relative uptake of the carrier by liver Kupffer cells and that this toxicity mechanism is dependent on the endosome disruptive function of the carrier. Finally, the PAF toxicity mechanism was shown to be generalizable to commercial delivery reagent *in vivo*-jetPEI^®^ and an MC3 lipid nanoparticle formulated to match an FDA-approved siRNA nanomedicine. Greater sensitivity to the PAF mechanism occurs in 4T1 tumor-bearing mice, a mammary tumor model known to exhibit increased circulating leukocytes and potential to respond to inflammatory insult. These results establish Kupffer cell release of PAF as a key mediator of *in vivo* nucleic acid nanocarrier toxicity and identify PAFR inhibition as an effective prophylactic strategy to increase maximum tolerated dose and reduce nanocarrier-associated adverse events.

**Significance:** Non-viral nucleic acid nanocarriers can enable *in vivo* gene therapy, but their potential interaction with innate immune cells can cause dose-limiting toxicities. Nanoparticle toxicities are currently poorly understood, making it difficult to identify relevant design criteria for maximizing nanoparticle safety. This work connects nanoparticle-associated toxicities to the release of platelet activating factor (PAF) by liver Kupffer cells. Small molecule inhibition of the PAF receptor (PAFR) completely prevents severe adverse events associated with high doses of multiple polymer-based formulations and a lipid nanoparticle matching the composition of the first clinically-approved siRNA nanomedicine. This study identifies PAF as a toxicity biomarker for future nanomedicine discovery programs. Further, PAFR inhibition should be explored as a strategy to expand the therapeutic index of nanomedicines.

## Introduction

With the recent first-in-class FDA approval of Onpattro in August 2018, nanoparticle-based delivery of siRNA has entered an era of clinical reality.(1) However, cationic carrier components continue to face toxicological challenges that hinder their clinical development. In fact, Alnylam Pharmaceuticals, the company that developed Onpattro, has abandoned its cationic DLin-MC3-DMA carrier lipids for their pipeline therapeutics due to the carrier-associated toxicities that necessitate immunosuppressive steroid pretreatment in patients.(2)

For cationic nanoparticle-based siRNA delivery systems (si-NPs), there is a correlation between transfection efficiency and toxicity which poses a challenge for maximizing the therapeutic window of siRNA nanocarriers.(3) Cationic polymeric carriers such as poly(ethylene imine) (PEI) have been studied extensively for their *in vitro* cytotoxicity mechanisms.(3-5) However, *in vivo* toxicity mechanisms are more complex and, as a result, not well-understood.(4) After intravenous injection, nanomaterials inevitably accumulate in cells of the mononuclear phagocyte system (MPS), especially in the liver.(6) Complement adsorption to nanoparticle surfaces can increase MPS clearance while also inducing inflammatory reactions.(3, 7) Aggregation of cationic si-NPs in the lungs can furthermore induce severe clotting resulting in acute, fatal toxicity in mice.(3, 8-10) Steric shielding of polyplexes with materials such as PEG is often considered a panacea to these toxicities.(3, 7, 10) However, PEGylated nanoparticles can still be toxic at high doses and cause poorly-understood allergic-type adverse reactions even in the absence of prior nanomaterial exposure (lacking IgE antibodies).(6, 11) CALAA-01, a PEGylated cyclodextrin-based siRNA carrier, was discontinued from clinical development in part due to infusion-related innate immune responses.(1, 12)

We previously developed novel siRNA-complexing polymers (PMPC-DB) containing a high molecular weight (20kDa) zwitterionic corona composed of a poly(2-methacryloyloxyethyl phosphorylcholine) (PMPC) homopolymer.(13) The core is composed of a poly(dimethylaminoethyl methacrylate-co-butyl methacrylate) (DMAEMA-co-BMA) 50:50 random copolymer which provides optimal endosomolytic behavior while reducing cytotoxicity.(14, 15) PMPC-DB si-NPs have improved circulation half-life compared to similar si-NPs with smaller molecular weight coronas and achieve greater tumor cell uptake compared to similar si-NPs with 20kDa PEG coronas.(13) When used at an ideal polymer:siRNA (N:P) formulation ratio and stabilized by “dual hydrophobization”, implemented by palmitate conjugation to the siRNA cargo, this si-NP is well-tolerated in long-term, multi-dose regimens.(16)

Despite this favorable profile, we have empirically observed that the zwitterionic si-NPs have notably narrower therapeutic window than the PEGylated si-NPs, manifested by acute toxicities in mice when intravenously delivered in a sub-optimal formulation (elevated dose, high N:P ratio, and absence of stabilizing hydrophobic functionalization of the siRNA). Mice intravenously treated under these conditions exhibit lethargy and prostrated body positions within 5-15 minutes and succumb within 1 hour of injection; gross pathology revealed congested livers, reddened intestines, and signs of vasodilation.

We hypothesized that these rapid, shock-like toxicities associated with particulate injection may be associated with platelet activating factor (PAF). PAF is a potent phospholipid signaling molecule that is produced by, and is an effector of macrophages, neutrophils, and platelets that triggers vasodilation, vascular permeability, and leukocyte degranulation.(17-21) PAF is associated with multiple systemic inflammatory responses, including sepsis and anaphylactic shock, and it has been associated with acute toxicities caused by intravenous adenovirus injection.(22, 23) PAF has never been linked to the toxicology of synthetic nanoparticles of any kind, despite the fact that nearly all intravenously-administered nanoparticles are internalized by Kupffer cells, specialized liver macrophages,(6, 24, 25) which are a rich source of PAF. (26) We therefore hypothesized that PAF secretion by Kupffer cells plays a heretofore unappreciated role in systemic non-viral nanoparticle toxicities.

## Results and Discussion

The PMPC-DB polymers used for this study were confirmed for structure by ^1^H-NMR (**Supplemental Figure S1**)(13) and were free of endotoxin contamination prior to initiation of all studies (**Supplemental Figure S2**). An si-NP formulation was prepared by complexing unmodified siRNA with PMPC-DB polymers at a 20:1 charge ratio (defined as protonated nitrogen to phosphate ratio), yielding monodisperse si-NPs with average hydrodynamic diameter of 100 (**Supplemental Figure S3**). This formulation was purposefully defined and used at a high dose (1.2 mg/kg siRNA, 73.6 mg/kg polymer), known to be poorly tolerated and to induce notable adverse events, in order to test the hypothesis that PAF is a key mechanism that drives si-NP toxicity.

We pre-treated BALB/c mice with either saline or ABT-491intravenously 10 minutes prior to injecting mice with si-NPs. ABT-491 is a potent and highly specific small-molecule inhibitor of the PAF receptor (PAFR).(21, 27) Of five mice pre-treated with saline, four mice experienced fatality within 1 hour of injection, (**Figure 1a**). Mice pre-treated with ABT-491 experienced no fatalities and no behavioral changes. Thirty minutes after si-NP injection, mice treated with si-NPs only (no PAFR inhibitor) exhibited gross pathology changes characteristic of shock and hypotension, including reddened intestines, congested livers, and vasodilation (**Figure 1b**). Mice pre-treated with ABT-491 did not experience these changes and appeared phenotypically similar to saline-treated mice.

**Figure 1.**
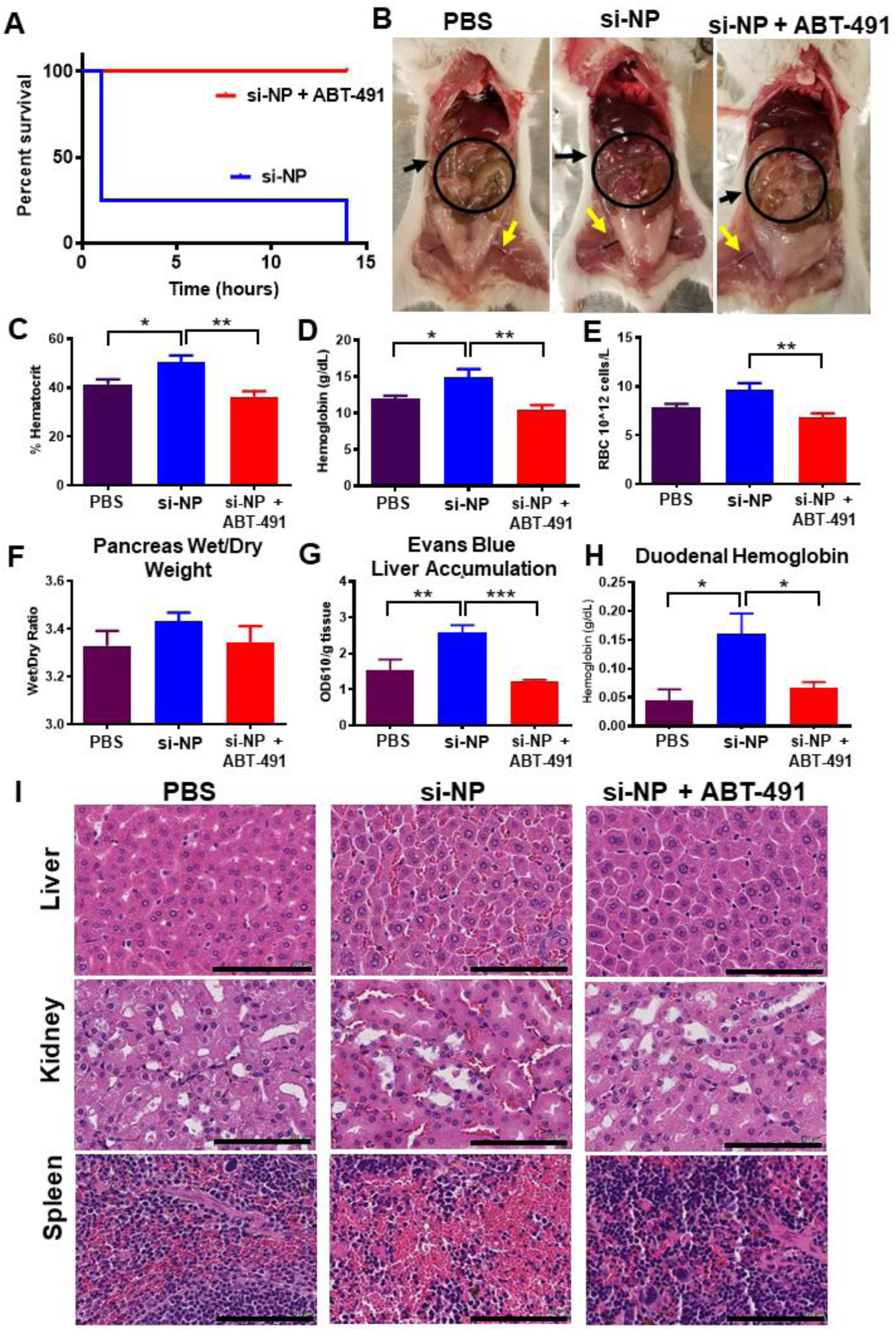
PAFR inhibitor ABT-491 rescues acute toxicities associated with high-dose si-NP injection. A) Upon intravenous injection of si-NPs at high N:P ratio (high polymer dose) and 1.2 mg/kg siRNA, 4 out of 5 mice experienced fatalities within 1 hour, but all deaths were prevented by pre-injection of ABT-491 10 minutes prior to si-NP injection (n=5 mice per group). B) Gross pathology demonstrates vasodilatory pathology in major blood vessels (yellow arrows) and reddening of intestines (black arrows, circles) in mice treated with si-NPs. C-E) Mice injected with high-dose si-NPs experienced significant increases in blood hematocrit (C), hemoglobin (D), and red blood cell concentration (E), all of which were abrogated by ABT-491 pretreatment. F) Pancreas wet weight vs. dry weight with 1.2 mg/kg si-NPs (n=5-7), saline, or si-NPs with ABT-491 pre-treatment. G) Evans Blue concentration in mouse livers. H) Duodenum hemoglobin concentration (n=5-7, * p<0.05, ** p<0.01, *** p<0.001). I) H&E staining of mouse liver, kidney, and spleen after i.v. injection demonstrating red blood cell congestion in each tissue. Scale bars = 100 µm. All measures were made 30 minutes after injection.

Mice treated with si-NPs experienced significant increases in blood hematocrit percentage, hemoglobin concentration, and red blood cell concentration at 30 minutes post-injection compared to saline-treated or ABT-491-pre-treated mice (**Figure 1c-e**). These blood hemoconcentration symptoms were indicative of shock. Shock is known to increase vascular permeability, which results in leakage of plasma fluid into tissues and consequent hemoconcentration. Hemoconcentration is also one of the hallmark side effects of intravenous PAF administration.(28)

Shock-induced vascular permeability can lead to edema in organs and GI tract effects, including increased duodenal hemoglobin.(22) We noted increases (though not statistically significant) in the pancreas wet/dry weight ratio for si-NP-treated mice compared to inhibitor-pre-treated mice (**Figure 1f**). We further confirmed these symptoms of tissue edema by injecting mice with Evans Blue dye five minutes after si-NP treatment and measuring dye absorbance in liver homogenates after si-NP treatment. Evans Blue dye binds to the serum protein albumin and can assess aberrant vascular permeability and plasma leakage into tissues.(29) Evans Blue dye absorbance was elevated in si-NP-treated mice compared to saline-treated mice, while ABT-491-pre-treated mice did not have increases (**Figure 1g**). These results indicate that ABT-491 pre-treatment rescued mice from si-NP-induced vascular permeability. si-NP-treated mice also had severe red blood cell congestion in their duodenums as indicated by nearly four-fold higher hemoglobin concentrations than saline-treated or ABT-491-pre-treated mice (**Figure 1h**).

Histologically, these results were confirmed by evidence of significant congestion in multiple organs of si-NP-treated mice (**Figure 1i**). Notably increased amounts of red blood cells were visible by H&E staining in the liver, kidney, and spleen of these mice. By comparison, mice pre-treated with ABT-491 did not experience histologic changes compared to saline-treated mice.

The combined data in Figure 1 suggests that PAF plays a significant role in the acute toxicities caused by high-dose si-NP injection. Inhibition of PAFR completely rescued mice from si-NP-associated toxicities such that there were no phenotypical or pathological differences observed between ABT-491+si-NP-treated mice and saline-treated mice. Additionally, we have demonstrated that the acute toxicities resulting from si-NP treatment are symptomatic of shock-like conditions including vasodilation, hemoconcentration, and vascular edema. These symptoms became apparent upon first exposure to si-NP treatments and are thus not consistent with anaphylactic, IgE-mediated shock. To our knowledge, this is the first confirmation of PAF involvement in nonviral nanoparticle-related toxicities.

The PAF receptor is expressed by a variety of immune cells, including neutrophils, eosinophils, macrophages, and platelets.(21, 23) Activation of PAFR by its ligand can induce neutrophil degranulation and/or inflammatory release of vasoactive agents such as leukotrienes or nitric oxide.(30) In macrophages, PAF is known to stimulate nitric oxide production which is believed to be the source of vascular permeability in PAF-mediated shock.(31-33)

We next sought to confirm whether the acute toxicities observed upon si-NP administration were dependent on mouse strain. Others have shown, for example, that mice strains that have Th-1 biased macrophages clear nanoparticles from circulation more slowly than Th-2-biased mice, leading to different pharmacokinetic and biodistribution profiles for the same nanoparticles in different mouse strains.(34) While BALB/c mice are generally more prone to Th-2 -type responses, C57BL/6 are Th-1 biased.(34) We therefore injected C57BL/6 mice with the same si-NP formulations and measured the effects of ABT-491 pre-treatment on blood hemoconcentration, pancreas edema, and duodenal hemoglobin (**Supplemental Figure S4**). Overall, we found that the shock-like toxicities seen with BALB/c mice were also consistently observed in C57BL/6 mice. Thus, the PAFR-mediated effects of si-NP administration are consistent across both Th-1 and Th-2 biased mouse strains.

To further understand the involvement of PAFR in si-NP-associated toxicities, we measured the blood PAF levels of si-NP-treated mice compared to saline-treated mice. Thirty minutes post-si-NP treatment, mice experienced significantly elevated blood PAF levels, ranging 6-8 ng/mL (**Figure 2a**). Because of the severe hemoconcentration caused by si-NP treatment, obtaining sufficient blood for PAF quantification was challenging, hindering our ability to acquire blood from the most severely affected mice. PAF can be lethal at doses as low as 0.36 µmol/kg, so it is reasonable that this PAF concentration would cause increased systemic toxicities.(28) These data indicate that intravenous administration of si-NPs induces systemic release of PAF.

**Figure 2.**
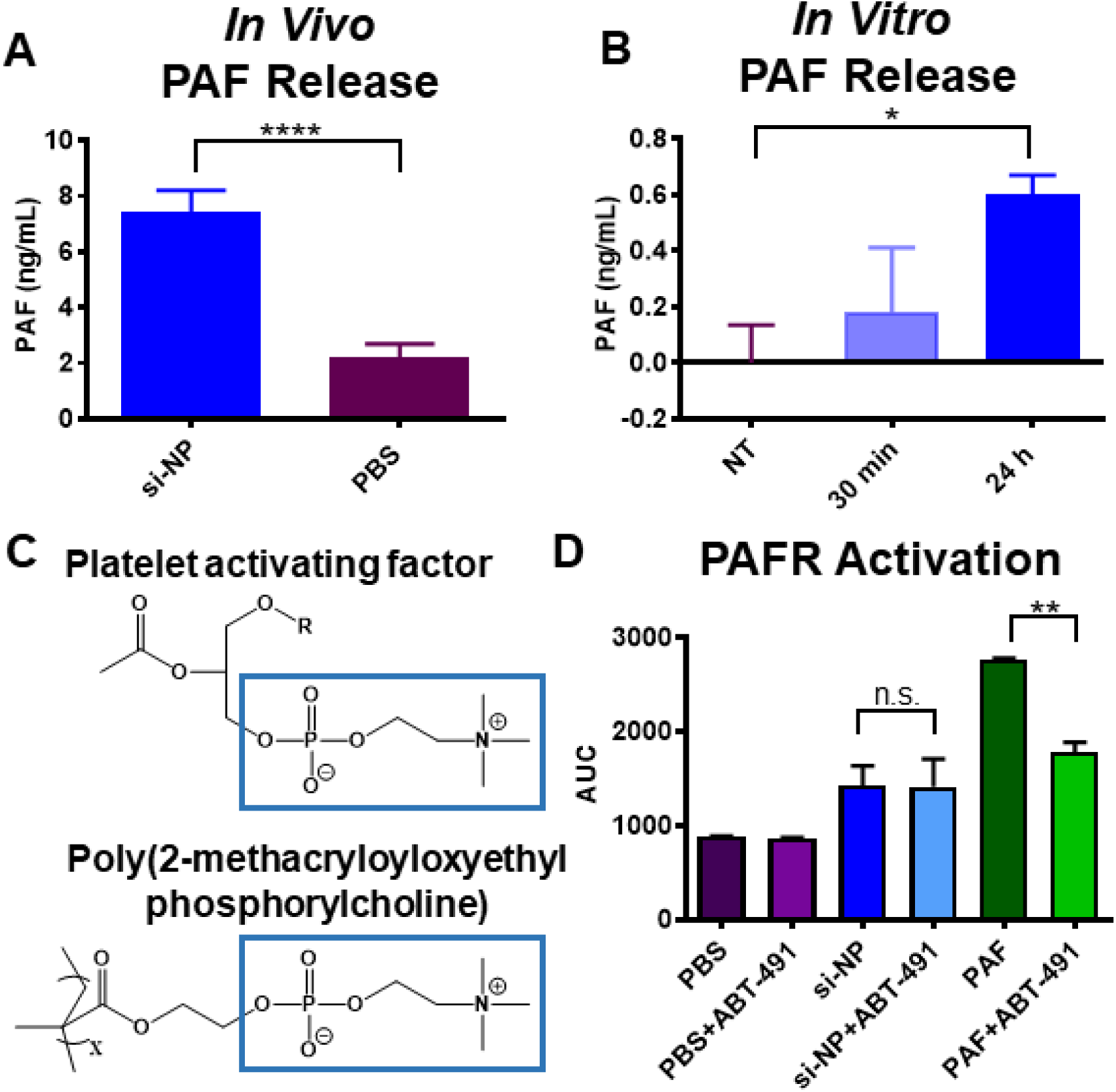
si-NPs stimulate PAF release and do not directly agonize the PAF receptor. A) Mice injected with si-NPs display increased PAF concentration in blood samples 30 minutes after injection. B) *In vitro*, mouse BMDMs stimulated with 100 nM si-NPs released PAF into culture media. C) The si-NP PMPC corona comprises zwitterionic phosphorylcholine, that are similar in structure to PAF. D) PAF activates the PAFR through a mechanism that is inhibited by ABT-491 pre-treatment in Chem-1 cells overexpressing PAFR, but PMPC-based si-NPs do not (* p<0.05, ** p<0.01, **** p<0.0001).

Because macrophages are one of the cell types known to release PAF, we stimulated mouse bone marrow-derived macrophages (BMDMs) with si-NPs for 24 hours and measured PAF release (**Figure 2b**). We found a significant increase in PAF in the media of si-NP-treated cells compared to saline-treated cells, suggesting that macrophages are one potential source of PAF release in si-NP-treated mice.

We furthermore hypothesized that our si-NPs themselves might be capable of agonizing the PAF receptor. The PMPC corona of these si-NPs is made up of zwitterionic, phosphorylcholine moieties, and the chemical structure of PAF itself contains a prominent phosphorylcholine (**Figure 2c**). Additionally, there have been examples in nature of bacteria using PAF-mimicry to engage the PAF receptor. For example, *Streptococcus pneumoniae* uses phosphorylcholine moieties in its cell wall to bind PAFR, block PAF signaling, and prevent inflammatory neutrophil reactions.(19, 35) We therefore treated Chem-1 cells overexpressing recombinant human PAFR with si-NPs and used an intracellular calcium dye to measure fluorescence from calcium influx (indicative of receptor activation). Addition of PAF itself caused a significant increase in cytosolic calcium that was reduced by pre-treatment with ABT-491. Addition of si-NPs caused a slight increase in fluorescent signal, but the effect was not reduced by ABT-491 pre-treatment (**Figure 2d**). These data suggest that, despite some chemical similarity, zwitterionic phosphorylcholine moieties on the si-NP surface do not directly act on the PAF receptor but instead stimulate systemic release of PAF, which then acts on PAFR to mediate vasoactive changes that cause si-NP toxicities. This effect is also consistent with studies showing that release of PAF causes dose-dependent increases in vascular permeability and hemoconcentration.(20, 28)

PAF can also be released when cells of the MPS are activated by phagocytosing material.(20) Nanoparticles in the 100 nm size range are well-known to be cleared from circulation by the MPS, particularly by liver Kupffer cells.(6) Kupffer cells reside intravascularly in liver sinusoids and serve as a phagocytic microbial filter for the blood, serving to remove nanoparticle and microbe-sized materials. To test whether Kupffer cells are responsible for si-NP treatment associated PAF release and consequent toxicities, we utilized clodronate liposomes to selectively deplete Kupffer cells. Clodronate liposomes are not toxic to other cell types, as they do not extravasate into tissues, and are only taken up by intravascular-residing phagocytic macrophages.(36)

Pre-treatment of mice with clodronate liposomes (Clod) prevented shock toxicities associated with intravenous si-NP injections (**Figure 3**). Clod-pre-treated mice receiving si-NPs did not experience significant elevations in hematocrit (**Figure 3a**), duodenal hemoglobin (**Figure 3b**), or blood PAF levels (**Figure 3c**), relative to Clod-treated mice receiving saline. These data indicate that macrophages of the MPS are primary mediators of PAF-related shock toxicities associated with si-NP injection. Our combined data indicate that Kupffer cells of the mouse liver, upon stimulation with intravenously delivered si-NPs, release PAF into the blood. PAF release activates downstream PAF-associated vasoactive signals and consequently causes hypotension, edema, and tissue congestion.

**Figure 3.**
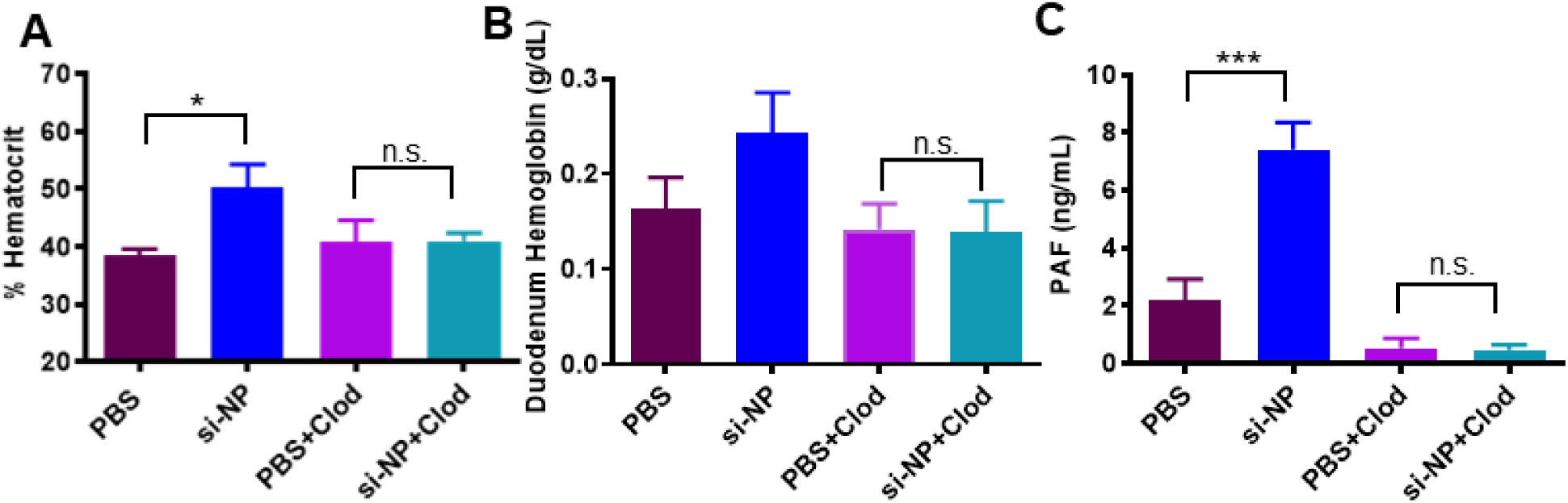
Pre-treatment with clodronate liposomes abrogates si-NP-related PAF elevation and toxicities. A) Mice pre-treated with Clod liposomes 24 hours prior to injection do not experience significant elevations in blood hematocrit 30 min after intravenous si-NP injection. B) Mice pre-treated with Clod liposomes do not experience significant increases in duodenal hemoglobin after intravenous si-NP injection. (n=4-5) C) Clod pre-treatment prevents increase in blood PAF levels after si-NP injection. Note that PBS and si-NP group data are reiterated from Figure 2a. (n=3-5, *p<0.05, ***p<0.001).

Kupffer cells provide a natural defense system for pathogenic, immunoreactive, nano-sized invaders, which can have similar features to nanoparticle drug delivery systems. PAF release from rat Kupffer cells has been associated with exposure to a variety of particular substances including adenovirus, zymosan particles, antibody-coated erythrocytes, immune complexes, and *Bordella pertussis*, so it is feasible that synthetic nanomaterials could also be a general trigger for PAF release.(37)

We hypothesized that if the PAF-related symptoms are nonspecifically initiated by Kupffer cell interactions with nanomaterials, then other polymeric nanoparticles would also stimulate PAF-related symptoms. In order to better understand the impact of si-NP structure on si-NP-associated toxicities, we compared our zwitterionic, phosphorylcholine surface si-NPs (zwitter si-NP) to si-NPs with a 20kDa PEG surface (matched molecular weight of PEG and PMPC polymer blocks) and the same endosomolytic DB core (PEG si-NP) (**Figure 4a**, polymer characteristics **Supplemental Figure S5A, S5C**). We also compared these si-NPs to a non-cationic, non-endosomolytic zwitterionic micelle (zwitter micelle) composed of a phosphorylcholine surface (PMPC) and a purely hydrophobic (BMA) core (**Figure 4a**, polymer characteristics **Supplemental Figure S5B, S5C**). These micelles were equivalent in size to both PEG si-NPs and zwitter si-NPs, but, due to their non-cationic nature, did not form complexes with siRNA (**Supplemental Figure S5D**).

**Figure 4.**
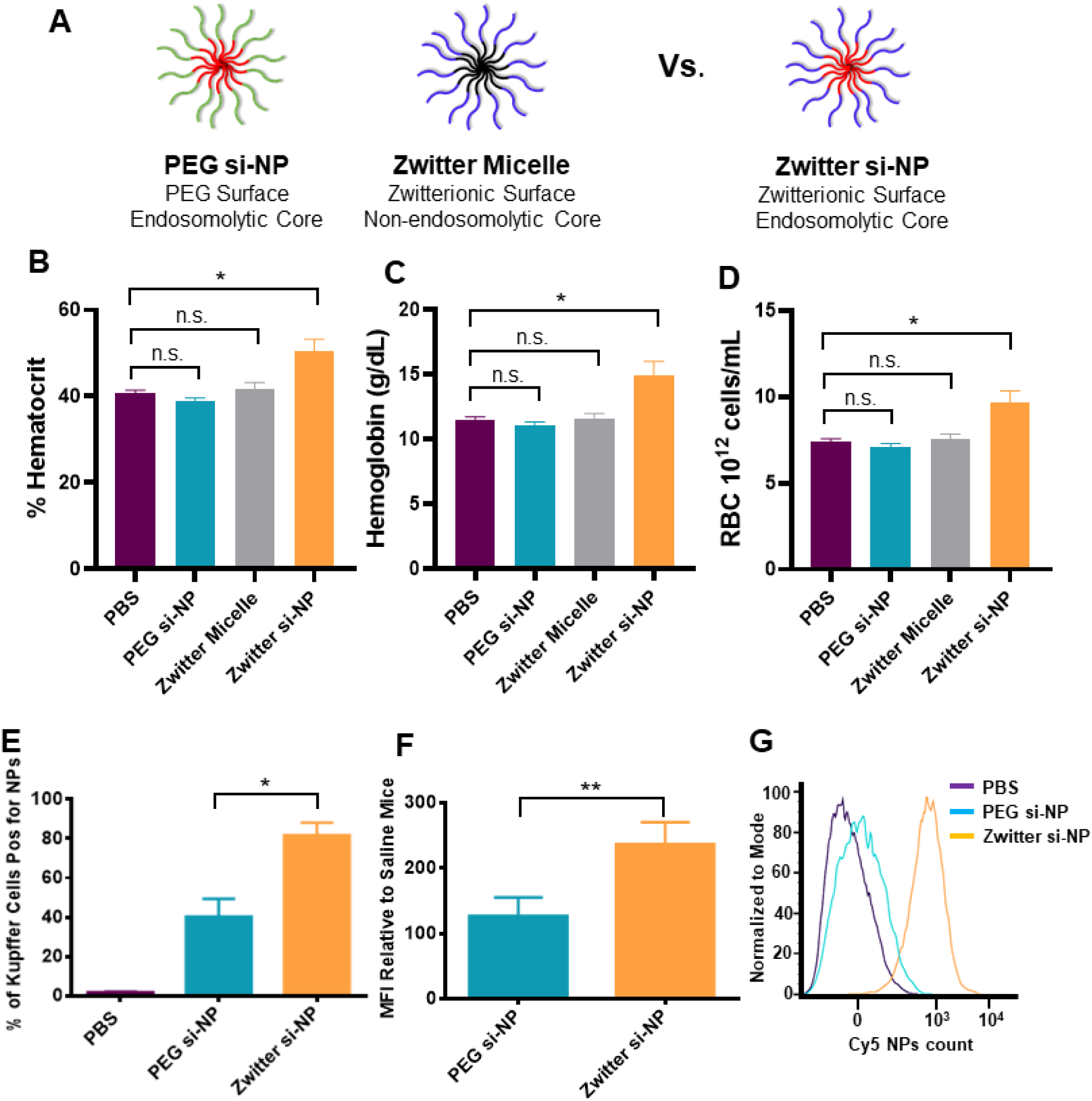
PAF-associated toxicities correlate with level of Kupffer cell delivery and are dependent on endosome-disruptive activity. A) Schematic of zwitter si-NP, PEG si-NP, and zwitter micelle structures tested. B-D) Hematocrit, hemoglobin, and red blood cell concentration values from mice 30 min after receiving a 1.2 mg/kg siRNA dose of each si-NP (n=3 mice per group, polymer dose: 73.6 mg/kg zwitter si-NP, 74.6 mg/kg PEG si-NP, 73.6 mg/kg zwitter micelle). E) Percent of mouse liver Kupffer cells positive for Cy5 si-NPs 20 min after intravenous administration. F) Mean fluorescence intensity (MFI) of PEG and zwitter si-NPs in mouse Kupffer cells relative to Kupffer cells of saline-treated mice (n=5, * p<0.05, ** p<0.01). G) Representative histograms of Kupffer cell uptake of fluorescent si-NPs after intravenous delivery.

In contrast to zwitter si-NPs, neither the PEG si-NPs nor the zwitter micelles caused changes in hematocrit, hemoglobin, or red blood cell concentration when given at an equivalent dose (**Figure 4b-d**). Similarly, PEG si-NPs or zwitter micelles did not cause increases in organ congestion (**Supplemental Figure S6**). These data suggest that the combination of a phosphorylcholine-based si-NP surface and an endosomolytic core may induce PAF-mediated shock, since the endosomolytic core (PEG si-NP) and phosphorylcholine corona (zwitter micelle) individually did not induce the same symptoms in mice when intravenously injected at an equivalent polymer dose. These collective data indicate that a combination of Kupffer cell uptake and endosome disruption is necessary for PAF-related toxicities. These findings are consistent with observations that innate immune activation by adenovirus can be prevented by impairing viral endosomal escape.(38)

To better understand the increased sensitivity to zwitter si-NPs compared to PEG si-NPs, we measured fluorescent si-NP uptake in Kupffer cells. In mice receiving injections of zwitter si-NPs, an average 82% of Kupffer cells took up si-NPs, while in mice receiving PEG si-NPs, only 40% of Kupffer cells took up the particles (**Figure 4e**). Additionally, mean fluorescent intensity was approximately two-fold higher in Kupffer cells from mice receiving zwitter si-NPs, indicating that this chemistry also achieved a higher amount of uptake per cell (**Figure 4f-g**) (Gating **Supplemental Figure S7**). These data suggest that si-NP PAF toxicities correlate to the level of acute delivery into Kupffer cells. We have previously shown that zwitter and PEG si-NPs do not adsorb complement proteins, have similar resistance to albumin adsorption, and also possess equivalent serum stability,(13) indicating that level of Kupffer cell internalization is the main driver of these toxicities. Increased cellular internalization of zwitter si-NPs by Kupffer cells is consistent with our previous observations that zwitter si-NPs were more highly internalized than PEG si-NPs by mouse tumor cells after intravenous injection.(13) Zwitter si-NPs suffer less from the “PEG dilemma” whereby increased PEGylation provides stability and improves systemic pharmacokinetics but comes at a significant cost to cell internalization and activity.(39) The increased uptake in zwitter si-NPs may result from differences in the physicochemical properties and molecular structures between PEG and zwitterionic PMPC. Water organizes in cage-like structures around PEG, but maintains its normal molecular organization around PMPC.(40) Additionally, the presence of phosphorylcholine moieties in PMPC may increase interactions with cell membranes, which contain phosphocholine-containing lipids.(41) Notably, we do anticipate that PEG si-NPs will be susceptible to PAF associated toxicities when delivered above their naturally maximum tolerated dose.

We next investigated whether PAF-associated toxicities are a generalizable dose-limiting mechanism that is relevant to other types of polymeric, pH-responsive nucleic acid delivery vehicles. These studies were also designed to examine whether systemic inflammatory status of the mice affects sensitivity to PAF-related toxicities, as a follow up to previous empirical observations that this IVJP dose causes toxicity in tumor-bearing mice.(42) We injected normal mice with a subtoxic dose of 2 mg/kg of a commercially available polymeric *in vivo* transfection reagent, *in vivo*-jetPEI^®^ (IVJP), which contains poly(ethylene imine) (PEI). PEI is considered a gold standard siRNA transfection reagent and is capable of triggering endosomolysis *via* the proton-sponge mechanism.(43) This dose did not reach the maximum tolerated dose in normal BALB/c mice (**Supplemental Figure S8**).

We completed parallel experiments with IVJP on mice bearing 4T1 mammary tumors. The metastatic 4T1 mammary tumor model induces a leukemoid reaction in mice, causing significant elevations in number of circulating neutrophils, monocytes, and other leukocytes (**Figure 5a-b**) through upregulation of 4T1 myeloid- and granulocytes-colony stimulating factors.(44) In human cancer patients, myeloid dysfunction and myelopoiesis is observed across many diverse tumor types, including breast, colon, pancreatic, prostate, and skin cancers.(45) Based on literature indicating that immune cells play a role in propagating and amplifying the vasoactive effects of PAF,(17-21) we hypothesized that PAF-associated toxicities would be enhanced in 4T1 tumor-bearing mice due to their increased number of circulating leukocytes.

**Figure 5.**
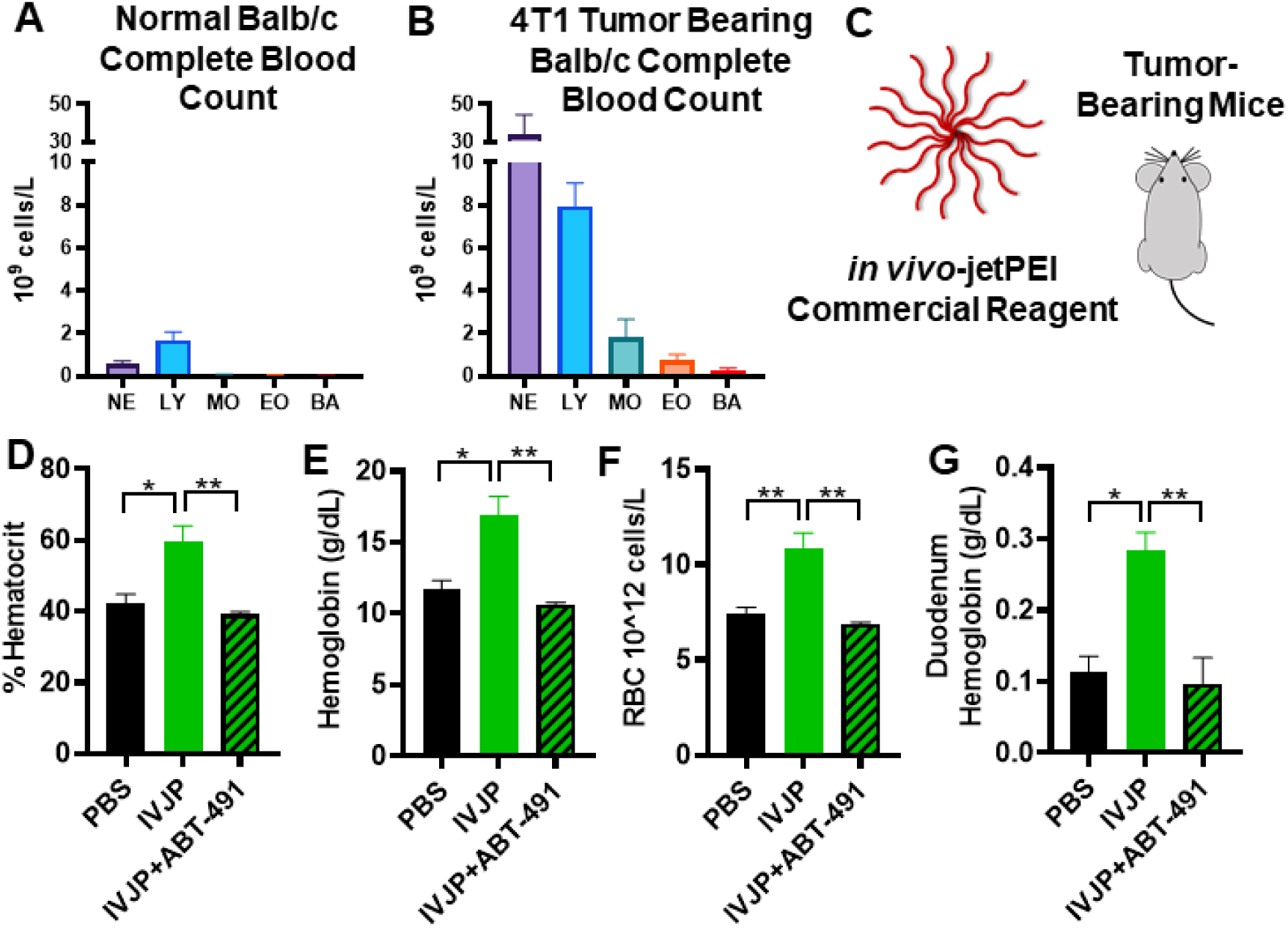
IVJP induces PAFR-dependent toxicity, which is enhanced in systemically-inflamed 4T1 tumor-bearing mice. A-B) Comparison of leukocytes in complete blood counts of untreated normal BALB/c mice or 4T1 tumor-bearing mice. C) Tumor-bearing mice were injected with 2 mg/kg IVJP D-G) Blood hematocrit, hemoglobin, red blood cell concentration, and duodenum hemoglobin concentration for tumor-bearing mice treated with saline, IVJP, or ABT-491 pre-treatment in combination with IVJP (n=3 mice per group, * p<0.05, ** p<0.01). NE, neutrophils; LY, lymphocytes; MO, monocytes; EO, eosinophils; BA, basophils.

In 4T1 tumor-bearing mice, IVJP treatment significantly increased blood hematocrit, hemoglobin, red blood cell concentration, and duodenum hemoglobin compared to saline-treated mice (**Figure 5c-g**), indicating PAF-induced toxicities in these mice. Pre-treatment with ABT-491 completely abrogated these blood changes, confirming involvement of PAFR in IVJP-related toxicity. Our data indicate that PAF-associated toxicities are generalizable to other types of pH-responsive, polymeric nanoparticle systems and that nanoparticle doses that are in a tolerable range for healthy animals can trigger PAF toxicities in scenarios where systemic tumor-associated inflammation is present.

We last sought to assess the generalizability of this mechanism to lipid-based systems. To do so, we tested a similar lipid nanoparticle (LNP) formulation to clinically-approved patisiran, including the ionizable lipid DLin-MC3-DMA (**Figure 6a**). While this LNP formulation is reported to be tolerated in non-tumor-bearing mice at doses as high as 15 mg/kg,(46) it induced rapid fatality in 4T1 tumor-bearing mice at 5 mg/kg (54.38 mg/kg DLin-MC3-DMA dose) (**Figure 6b**). Pre-treatment of these mice with the PAFR inhibitor ABT-491 completely prevented fatalities (**Figure 6b**). Complete blood count measurements taken 20 minutes after LNP injection revealed increased hematocrit, hemoglobin, and red blood cell concentrations (**Figure 6c-e**). Mice pre-treated with ABT-491 prior to LNP injection did not experience significant increases in hemoconcentration compared to saline-treated mice. These data indicate that inhibition of the PAF receptor prevents LNP-induced toxicities in tumor-bearing mice and suggest that PAF plays a role in not only polymer nanoparticle-associated toxicities but also LNP-associated toxicities. This finding has broad clinical implications, as patisiran and other LNP-based formulations are either currently FDA-approved or in clinical trials. Particularly for LNP formulations being used to treat cancer patients, the role of PAF in injection-related adverse events should be explored further.

**Figure 6.**
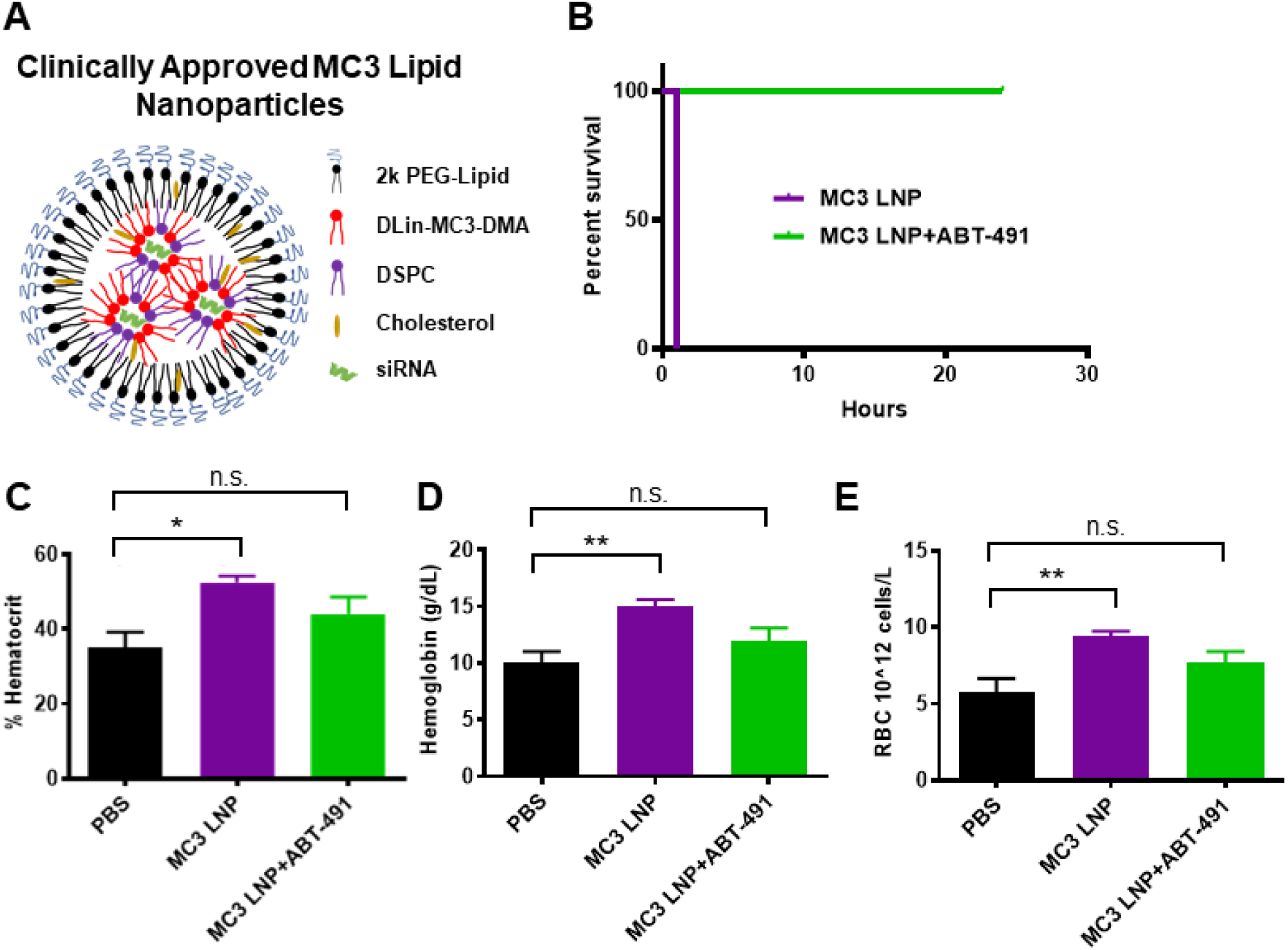
PAF-related toxicities are generalizable to lipid nanoparticles. A) Schematic illustrating components of patisiran LNP formulation. B) Survival curve for mice injected intravenously with 5 mg/kg MC3 LNP formulation with either saline or ABT-491 pre-treatment. (n=2 mice per group) C-E) Hematocrit, hemoglobin, and red blood cell concentration values of mice treated with saline, MC3 LNPs, or MC3 LNPs with ABT-491 pre-treatment (n=4-5, * p<0.05, **p<0.01)). All blood measurements were made 20 min after administration of MC3 LNPs.

Sensitivity to PAF-related toxicities correlated in both of these studies with the number of circulating leukocytes. These results are consistent with previous data showing that 4T1 tumors sensitize mice to anaphylactoid-type reactions in response to adenovirus treatment.(47) The increased sensitivity to PAF signaling in 4T1 tumor-bearing mice was associated with the presence of increased numbers of circulating leukocytes, highlighted by a 22-fold increase in monocytes and a 61-fold expansion of circulating neutrophils. PAF is known to activate monocyte secretion of inflammatory cytokines and to promote chemoattraction, adhesion, and degranulation of neutrophils, all of which can affect vascular permeability(48-52). Our collective data indicate that Kupffer cells are stimulated by high doses of nucleic acid nanomedicines to release PAF, which in turn acts through circulating leukocytes as an intermediate that produces vasoactive agents (such as nitric oxide)(31-33, 53) and consequent shock-like toxicities. (**Figure 7**).

**Figure 7.**
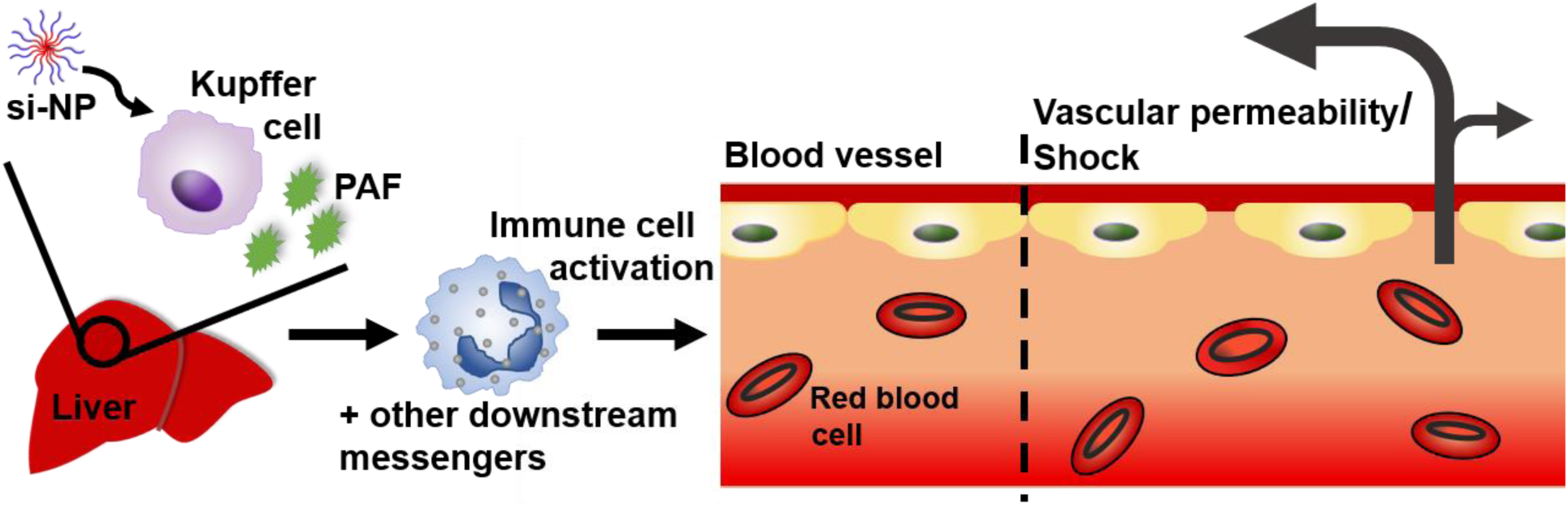
Schematic of proposed mechanism for PAF-mediated effects upon intravenous administration of nucleic acid nanocarriers.

Our findings suggest important implications particularly for the use of nucleic acid delivery vehicles in cancer treatment. The increased sensitivity of tumor-bearing to PAF-mediated toxicity may carry over to human cancer patients, who often exhibit myeloid dysfunction, elevated platelet counts, and an increased risk of clotting due to the secretion of pro-clotting factors by tumor macrophages.(44, 45, 54) While PAFR is present on all of the same cell types in humans as in rodents, humans additionally possess PAFR on platelets.(17) Thus, nanoparticle and PAF-mediated toxicities are expected to be similar or potentially even more pronounced in humans, particularly in cancer patients who are in a hyper-coagulative state. Indeed, adverse events reported in the Phase 1a/1b clinical trial for cancer patients receiving cyclodextrin-based siRNA carrier CALAA-01 included hypersensitivity and GI effects such as ischemic colitis and edema as well as events such as thrombocytopenia and tachycardia, all of which can be related to effects caused by PAF.(12) There are additional reports of hypotension-related adverse effects in cancer patients injected with adenovirus vectors.(22)

While no studies have yet connected nanoparticle toxicities to PAF, there have been multiple reports of nanocarriers inducing toxicities that may have unknowingly been related to PAF in both rodents and humans. For example, multiple studies in rodents have reported rapid fatalities within the first hour of injection of PEI polyplexes, although this lethality is often blamed on pulmonary clotting due to cationic polymer components.(9, 10) However, other in-depth studies of *in vivo* PEI-mediated toxicities have shown evidence of liver necrosis, shock, and smaller aggregates of CD11b+ cells and platelets, without major lung obstruction.(8) Other types of non-cationic nanocarriers have also been shown to induce poorly-understood cardiovascular changes in mice, including carbon nanotubes, acrylic polymers, and mesoporous silicon.(55, 56) In humans, adenovirus and nanoparticle Doxil injections have been associated with adverse events including hypotension and decreased cardiac output.(57, 58) Our work suggests that the role of PAF in the response to intravenous nanocarriers should be further explored for other types of nanomaterials.

Our results suggest that inhibition of PAFR may help prevent intravenous nanomedicine-associated toxicities. Prophylactic treatment protocols are already the norm in the clinical translation of nanomedicine---Onpattro, the only FDA-approved siRNA nanomedicine, requires pre-treatment of patients with corticosteroids to dampen immune activation.(1) While no PAFR antagonists have yet been clinically approved in the U.S., rupatadine is a combination anti-histamine and PAFR inhibitor approved for use in other countries.(18, 22) Our results suggest that PAFR inhibitors such as ABT-491 or rupatadine may find new clinical benefit by making nanomedicines safer and to potentially enable use of higher, more efficacious doses by expanding their therapeutic index.

## Conclusion

Overall, this work provides compelling evidence that PAF is the key factor in determining dose-limiting toxicities of polymer- and lipid-based siRNA nanomedicines. To our knowledge, PAF has never been connected to synthetic, non-viral nanocarrier toxicities associated with intravenous injection. We have shown that PAF release is dependent on nanocarrier interaction with Kupffer cells and that the presence of systemic inflammation, associated with higher circulating immune cells, predisposes mice to PAF-mediated toxicities. We have also shown that PAF-related toxicities are generalizable to other types of polymeric and lipid nanocarriers, indicating that this may be an unrealized and under-studied toxicity mechanism for other types of particles. Inhibition of PAFR prior to nanoparticle administration can completely prevent acute shock-associated toxicities in mice, suggesting that PAFR is a viable prophylactic target that can be used to broaden the therapeutic window and improve clinical management of adverse events related to siRNA nanomedicines.

## Materials and Methods

All materials were purchased from Sigma-Aldrich (St. Louis, MO, USA) or Fisher Scientific (Waltham, MA, USA) unless indicated otherwise. siRNAs used in these studies were purchased from Integrated DNA Technologies (Coralville, IA, USA) and consisted of a sense strand (5’-CAAUUGCACUGAUAAUGAACUCC[dT][dC]-3’) and antisense strand (5’-GAGGAGUUCA[mU]U[mA]A[mU]U[mG][mU][mU]-3’) [Note: m=2’O-methyl modification; d=chimeric DNA base]. In flow cytometry experiments, Cy-5 labeled DNA oligonucleotides (IDT) were used instead of siRNA, and consisted of sense strand (5’-dGdTdCdAdGdAdAdAdTdAdGdAdAdAdCdTdGdGdTdCdAdTdC-3’) and antisense strand (5’-[Cy5]dGdAdTdGdAdCdCdAdGdTdTdTdCdTdAdTdTdTdCdTdGdAdC-3’).

### *Polymer Synthesis and Characterization* (NMR, GPC, DLS)

Polymers were made using Reversible Addition Fragmentation Chain Transfer (RAFT) polymerization using a chain transfer agent (CTA) of 4-(ethylsulfanylthiocarbonyl)sulfanylpentanoic acid (ECT). All monomers except 2-methacryloyloxyethyl phosphorylcholine were passed twice through activated alumina columns to remove inhibitors. PMPC-DB polymers (poly(2-methacryloyloxyethyl phosphorylcholine (PMPC))-*b*-(dimethylaminoethyl methacrylate (DMAEMA)-*co*-butyl methacrylate (BMA))) were synthesized by first creating a random copolymer of 50% DMAEMA and 50% BMA monomers polymerized from ECT in dioxane under nitrogen (30-minute purge) for 24 h at 65°C using azobisisobutyronitrile (AIBN) as an initiator (1:10 CTA:AIBN ratio). Equivalent molar amounts of DMAEMA and BMA were added to target an overall degree of polymerization (DP) of 210 repeating units overall (anticipating ∼ 70% conversion and final DP of 150-160). The crude reaction was precipitated in ice cold pentane three times to remove monomers and dried under vacuum overnight. DP was confirmed using ^1^H-NMR and GPC. From this purified first block (DB-ECT), the PMPC hydrophilic, si-NP surface-forming block was polymerized in methanol for 24 hours at 65°C under nitrogen (30-minute purge) with a 1:5 CTA:AIBN ratio. A DP of 75 was targeted to achieve an approximately 20,000 g/mol corona block (anticipating 90-95% conversion). The crude reaction mixture was then dialyzed overnight in methanol and then in deionized water for one day. The polymer was lyophilized, and PMPC block degree of polymerization was confirmed with ^1^H-NMR. The control PMPC-BMA polymer (zwitter micelle) was similarly synthesized, starting from a PBMA homopolymer. The PMPC block was polymerized in pure ethanol with all other conditions the same to improve solubility of the PBMA macroCTA. The 20kPEG DB polymer (PEG si-NP) was synthesized and characterized as described previously.(13) All NMR spectra are featured in **Supplemental Figures S1, S5A-B, S9**. Polymers were evaluated for endotoxin using a Chromogenic LAL Endotoxin Assay Kit (GenScript, Nanjing, China).

### si-NP Formulations

Zwitter si-NPs and PEG si-NPs were formulated as described previously.(13) Briefly, polymers were complexed with siRNA at a 20:1 N:P ratio (protonated nitrogens on polymer: anionic phosphates on siRNA) in 10 mM citrate buffer at pH 4. Polymers were initially dissolved at 30 mg/mL in pure ethanol and then diluted to 3 mg/mL in the pH 4 citrate buffer. After 30 minutes, samples were supplemented by addition of a 5x volume excess of 10 mM phosphate buffer at pH 8 to achieve a final pH of 7.4. PMPC-BMA micelles were dissolved in ethanol (5% of the final si-NP solution volume) and added dropwise to PBS for a final concentration equivalent to the polymer concentration of PMPC-DB for delivering 1.2 mg/kg siRNA (roughly 18 mg/mL). For *in vivo* preparations, polyplexes and micelles were sterile filtered (0.45 µm filters) and then concentrated to the appropriate volume (1.2 mg/kg dose in 100 μL injections) in Amicon spin filters (MWCO 50,000 kDa).

### PAFR Inhibition Studies

Mice (BALB/c, C57BL/6) were purchased from Jackson Laboratories (Bar Harbor, ME) or Charles River Laboratories (Wilmington, MA). For PAFR inhibition studies, mice were pre-injected with either 50 µL saline or 50 µL of ABT-491 (0.05 mg/mL in PBS -/-). Ten minutes after the first injection, mice were then injected with 1.2 mg/kg si-NPs (PMPC-DB polyplexes). A separate cohort was injected with saline only. For hematology and pathology studies, mice were sacrificed 30 minutes following si-NP injection, and blood was harvested by cardiac puncture using EDTA as an anti-coagulant. Blood samples were analyzed for complete blood counts (including hematocrit, hemoglobin, and red blood cell measures) by the Vanderbilt Translational Pathology Shared Resource using a Forcyte Analyzer (Oxford Science, CT, USA). Mouse organs were removed, except for the pancreas and duodenum, and were fixed in 10% formalin for histologic analysis.

The pancreas was weighed after removal, frozen, lyophilized, and weighed again for pancreas wet/dry weight analysis. The duodenum was washed with a saline solution containing 0.16 mg/mL heparin to remove excess blood. The tissue was then weighed and homogenized in 1 mL of saline/heparin solution. The tissue was centrifuged at 10,000 xg for 15 minutes at 4°C. Supernatants were removed and assayed for hemoglobin content using a Hemoglobin Colorimetric Assay Kit (Cayman Chemical No 700540) according to manufacturer instructions. Fixed tissues were embedded in paraffin and stained with hematoxylin and eosin by the Vanderbilt Translational Pathology Shared Resource. For Evans Blue measurements of vascular leakiness, mice were injected with 50 µL of a 0.5% (wt/v%) Evans Blue solution in saline that had been sterile-filtered through a 0.45 µm filter 2 minutes after si-NP (or saline) injection. After sacrificing mice at 30 minutes post-injection, livers were removed, weighed, and then incubated in 1 mL formamide for 24 hours at room temperature. A 50 uL aliquot of formamide for each sample was then transferred to a 96-well plate, and Evans Blue absorbance was measured at 610 nm using a plate reader (Tecan, Mannedorf, Switzerland).

### Clodronate Liposome Studies

Clodronate liposome pre-treatment was used to deplete Kupffer cells.(59) For animal studies involving clodronate pre-treatment, half of the mice were injected both i.v. and i.p. with 100 µL of Clodronate Liposomes (Cedarlane, Burlington, Canada) 24 hours prior to si-NP treatments. Both normal mice and clodronate-treated mice were then either injected with si-NPs or saline 24 hours later and sacrificed 30 minutes after si-NP injection. Organs and blood were collected and processed as described above.

### PAF ELISA

Mouse EDTA-treated blood samples were centrifuged at 3,000xg for 10 min at 4°C and plasma was isolated for ELISA analysis. Mouse PAF ELISA kits were purchased from LifeSpan Biosciences (Seattle, WA, USA). Plasma samples were assayed in duplicate according to kit manual using 50 µL samples in each well. For *in vitro* PAF release assays, mouse bone marrow-derived macrophages (BMDMs) were extracted from the femurs and tibias of healthy female FVB mice sacrificed at 4-8 weeks of age. Bone marrow was flushed out with DMEM using a 5 mL syringe and collected in DMEM on ice. The cell suspension was centrifuged at 1000xg for 5 min and the pellet was resuspended in 2 mL ACK Lysis buffer and incubated for 2 minutes on ice. The lysis solution was then diluted in 20 mL warm DMEM (37°C) and centrifuged again at 1000xg for 5 minutes. BMDMs were then re-suspended in 10 mL of BMDM media (DMEM, 10% FBS, 1% PenStrep, 1% L-glutamine, 14% (1:1 v/v) L929 media week 1 and week 2 media [created by culturing L929 murine fibroblast cells in complete DMEM and collecting media after 7 days of incubation [week 1] and again 7 days later [week 2]]). The cells were then seeded in a 96 well plate at 1×10^5^ cells per well. BMDMs were polarized to an M1 phenotype by incubation with M1-inducing cytokines.(60) Briefly, 7 days after seeding (after replenishing media on day 2 and day 4), M1 BMDMs received media supplemented with 0.1 µg/mL IFN-γ and 0.1 ng/mL LPS. 24 hours later, media was removed, cells were washed, and 100 nM si-NPs were introduced to cells. Cell supernatants were sampled at 30 minutes and 24 hours post-treatment and assayed using mouse PAF ELISA kits.

### PAFR Activation Studies

Ready-to-Assay Platelet Activating Factor Receptor Cells (Millipore Sigma, Burlington, MA) (Chem-1 host cells overexpressing PAFR GPCR) were thawed and seeded at 14,000 cells per well in a 384-well plate according to manufacturer instructions. Cells were then washed with HBSS (20 mM HEPES) and loaded with FLIPR Calcium 6 dye (dissolved in HBSS [20mM HEPES, 2.5 mM probenecid] and incubated for 2 hours according to FLIPR Calcium 6 kit instructions. si-NPs were prepared at 50 nM siRNA (final in-well concentrations) in a 384-well plate, and ABT-491 was prepared at 135 μM in a separate plate. Fluorescence from calcium influx was assayed using the Vanderbilt High Throughput Screening Facility, on a Panoptic instrument (WaveFront Biosciences, Franklin, TN). Five µL of ABT-491 or saline were first added to the cells, incubated for 5 minutes, and then 12.5 µL of si-NP preparations, saline, or PAF positive control (28.3 μg/mL stock) were added to the cells, and fluorescence was recorded once per second for 20 minutes following si-NP addition.

### In Vivo *Kupffer Cell Uptake*

Wild type BALB/c mice (Jackson Laboratories) were injected with either saline or with zwitter si-NPs or PEG si-NPs bearing 1.2 mg/kg Cy5-labeled DNA, prepared as described above. Twenty minutes after injection, mice were euthanized, perfused with PBS, and livers were removed and placed in 3 mL RPMI containing 5% FBS. An additional 3 mL of RPMI containing 4 mg/mL Collagenase II was added and incubated at 37°C for 30 minutes on an orbital shaker at 200 rpm. After digest, livers were filtered through 100 μm filters and hepatocytes were pelleted at 50xg for 2 min. Supernatants were collected and spun at 1200xg for 8 minutes. Pelleted cells were resuspended in 10 mL of clear 1X HBSS. A Percoll gradient was prepared consisting of a bottom layer of 20 mL 50% Percoll and a middle layer of 20 mL of 25% Percoll. Cells in HBSS suspension were then gently added to the top of the gradient. Samples were centrifuged 15 min at 800xg with no brakes. The buffy coat interface layer was removed carefully and placed in 40 mL of PBS containing 1% FBS. Cells were centrifuged 5 min at 500xg and then subjected to red blood cell lysis in 3 mL ACK buffer, after which time 20 mL of PBS (1% FBS) were added, and cells were again spun at 500xg for 5 min. Cells were resuspended in 400 uL FACS buffer containing Fc block (1:100) for 5 minutes, and passed through a 35 μm filter. Cells were centrifuged again at 500 xg for 5 minutes, and then resuspended in 200 uL FACS buffer. Samples were kept on ice.

Cells were immunolabeled using antibodies (Biolegend) reactive against F4/80 (PE; Clone BM8; 1:200), CD45 (PerCp-Cy5.5; Clone 30-F11; 1:4000), and CD11b (FITC; Clone M1/70; 1:200). Immediately prior to flow cytometry, cells were stained for viability with 4’,6-diamidino-2-phenylindole (DAPI, 0.25 μg/mL). Flow cytometry was performed on a MACSQuant Analyzer 10 (Miltenyi Biotec). Data analysis was performed using FlowJo software (FlowJo LLC, Ashland, OR). Kupffer cell gating was performed as described in Lynch et al and absence of non-specific antibody binding was confirmed using fluorescence-minus-one controls.(61)

### Tumor Mice, IVJP, and LNPs

Seven-week-old female BALB/c mice were implanted with 1×10^5^ 4T1 cells per tumor (in 50:50 mixture of serum-free DMEM and Matrigel (Corning, #354234) in the inguinal mammary fatpad (2 tumors per mouse). Nanoparticle and micelle treatments were initiated once tumors had reached 150-300 mm^3^. All tissue processing/endpoint timing occurred as stated above for the PAFR inhibition studies. *In vivo*-jetPEI (IVJP) (Polyplus, NY, USA) was prepared according to manufacturer instructions, at a dose of 2 mg/kg siRNA. Lipid nanoparticles (LNPs) were prepared at a dose of 5 mg/kg siRNA, and consisted of DLin-MC3-DMA [(6Z,9Z, 28Z, 31Z)-Heptatriaconta-6,9,28,31-tetraen-19-yl 4-(dimethylamino) butanoate), (MedKoo Biosciences, Morrisville, NC)], DSPC (1,2-distearoyl-sn-glycero-3-phosphocholine, Avanti Polar Lipids Inc, AL, USA), Cholesterol (Avanti Polar Lipids Inc, AL, USA), and DMG-PEG 2000 (1,2-dimyristoyl-rac-glycero-3-methoxypolyethylene glycol-2000, Avanti), prepared at molar ratios of 50:10:37.5:2.5 and a total siRNA:lipid weight ratio of 0.06, as described in Yamamoto et al.(62) Lipids and siRNA were mixed in a NanoAssemblr (Precision NanoSystems, Vancouver, BC, CA), at a flow rate of 2 mL/min (1:2 flow ratio, lipids dissolved in ethanol at 33% of final volume, siRNA in sodium citrate buffer, pH 4). LNP formulation was dialyzed in sterile PBS -/- overnight prior to i.v. administration. Mice injected with LNPs were sacrificed 20 minutes after injection.

## Author Contributions

M.A.J, S.S.P, T.D.G, and C.L.D. conceived and designed the research. M.A.J. and S.S.P synthesized the polymers and nanoparticles and performed the *in vitro* studies. M.A.J, S.S.P, E.N.H, I.B.K, S.K.B, A.R.K, R.E.M, D.D.L, P.P assisted in normal animal and tumor PAF studies. F.Y. performed blood isolation. B.R.D assisted in duodenum isolation and quantification. E.B.G performed BMDM isolation for *in vitro* PAF release. M.A.C and A.M.H assisted in Kupffer cell flow cytometry experiments. M.A.J, S.S.P, and C.L.D wrote the manuscript with feedback from all the authors.

## Acknowledgements

This work was supported by the DOD (DOD CDMRP OR130302), NIH (NIH R01 CA224241, NIH R01 EB019409, NIH R01 DK084246), NSF CAREER BMAT 1349604, the Vanderbilt Engineering and Immunity Pilot and Feasibility Grant, and the National Science Foundation (NSF GRF 1445197, NSF GRF 1937963).

The authors also acknowledge the assistance of the Vanderbilt Translational Pathology Shared Resource (TPSR), supported by NCI/NIH Cancer Center Support Grant 2P30 CA068485-14 and the Vanderbilt Mouse Metabolic Phenotyping Center Grant 5U24DK059637-13, for serum chemistry and complete blood counts.

