## Supplemental Information for "Kupffer Cell Release of Platelet Activating Factor Drives Dose Limiting Toxicities of Nucleic Acid Nanocarriers"

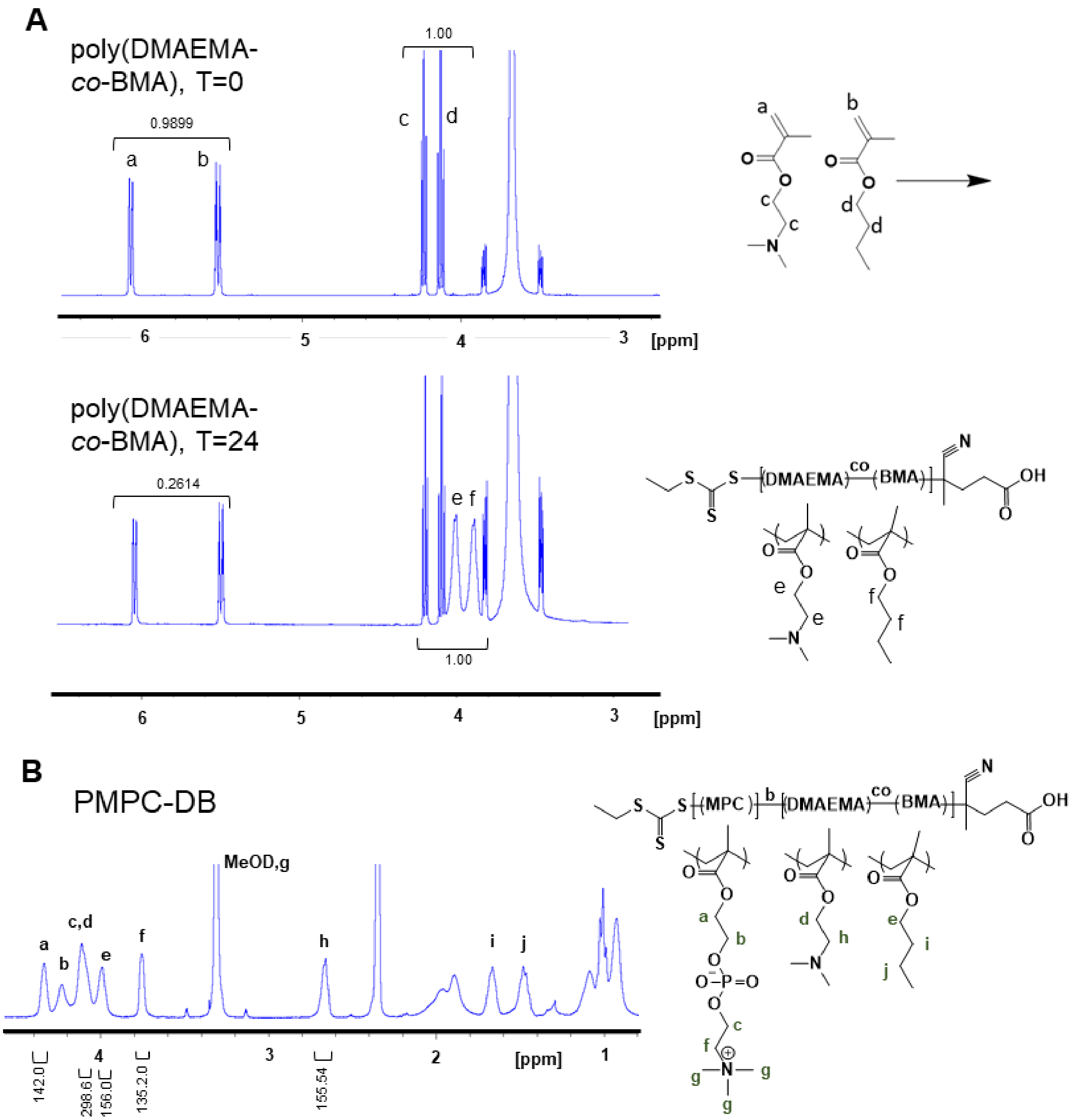


**Supplemental Figure S1**: A) [1]H-NMR of synthesis of poly(DMAEMA-co-BMA). Monomer conversion was calculated based on the disappearance of monomer peaks, as shown ((0.9899-0.2614)/0.9899), CDCl3 solvent. B) [1]H-NMR characterization of PMPC-DB, MeOD solvent.

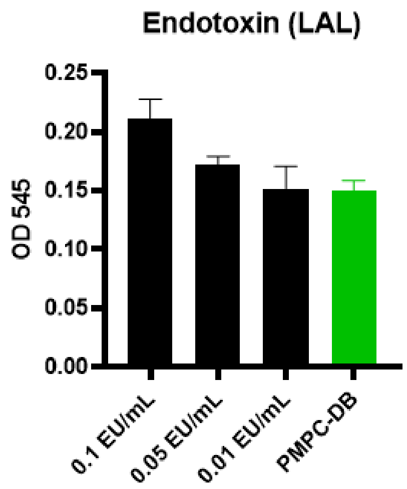


**Supplemental Figure S2**: PMPC-DB polymers were free of endotoxin contamination, as determined by the chromogenic LAL test (GenScript).

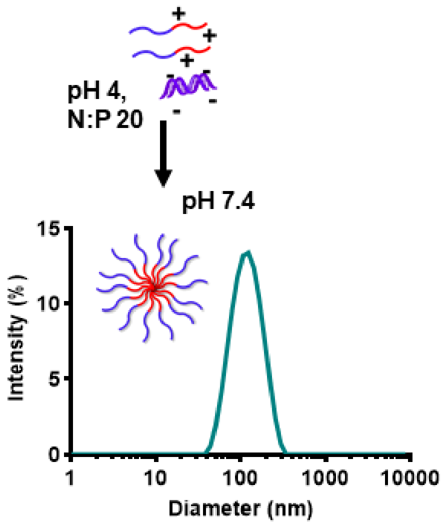


**Supplemental Figure S3**: si-NPs are formulated at N:P 20 by complexing with siRNA (non-hydrophobically modified) at a reduced pH. Raising pH to 7.4 produces si-NPs approximately 100 nm in hydrodynamic diameter.

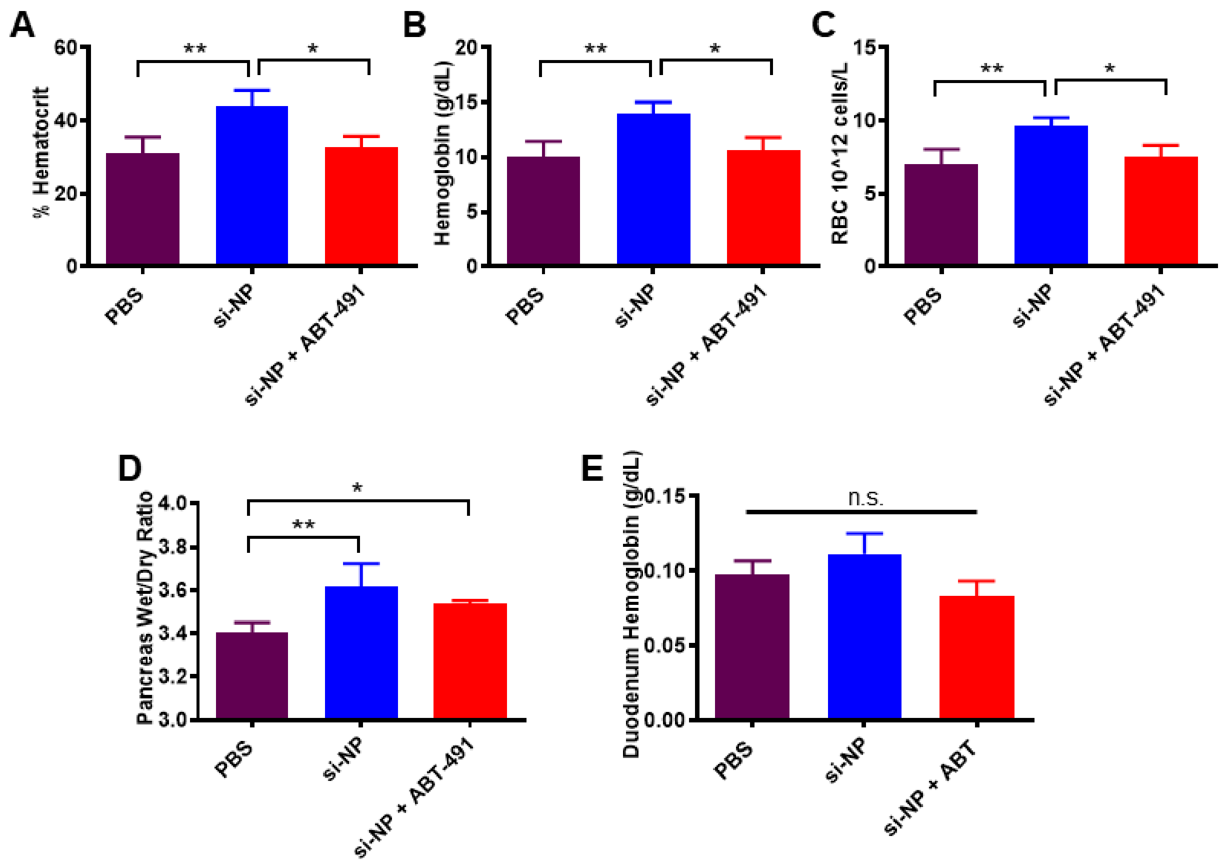


**Supplemental Figure S4**. Vasodilatory and edema-related toxicities and impact of PAFR inhibition are consistent in the C57BL/6 strain with initial observations in the BALB/c strain. A-C) C57BL/6 mouse hematocrit, hemoglobin, and red blood cell concentration 30 minutes after injection of si-NPs, saline, or si-NPs with ABT-491 pre-injection. D) Pancreas wet/dry ratio, E) duodenum hemoglobin concentration for the same mice (n=5, * $p<0.05$).

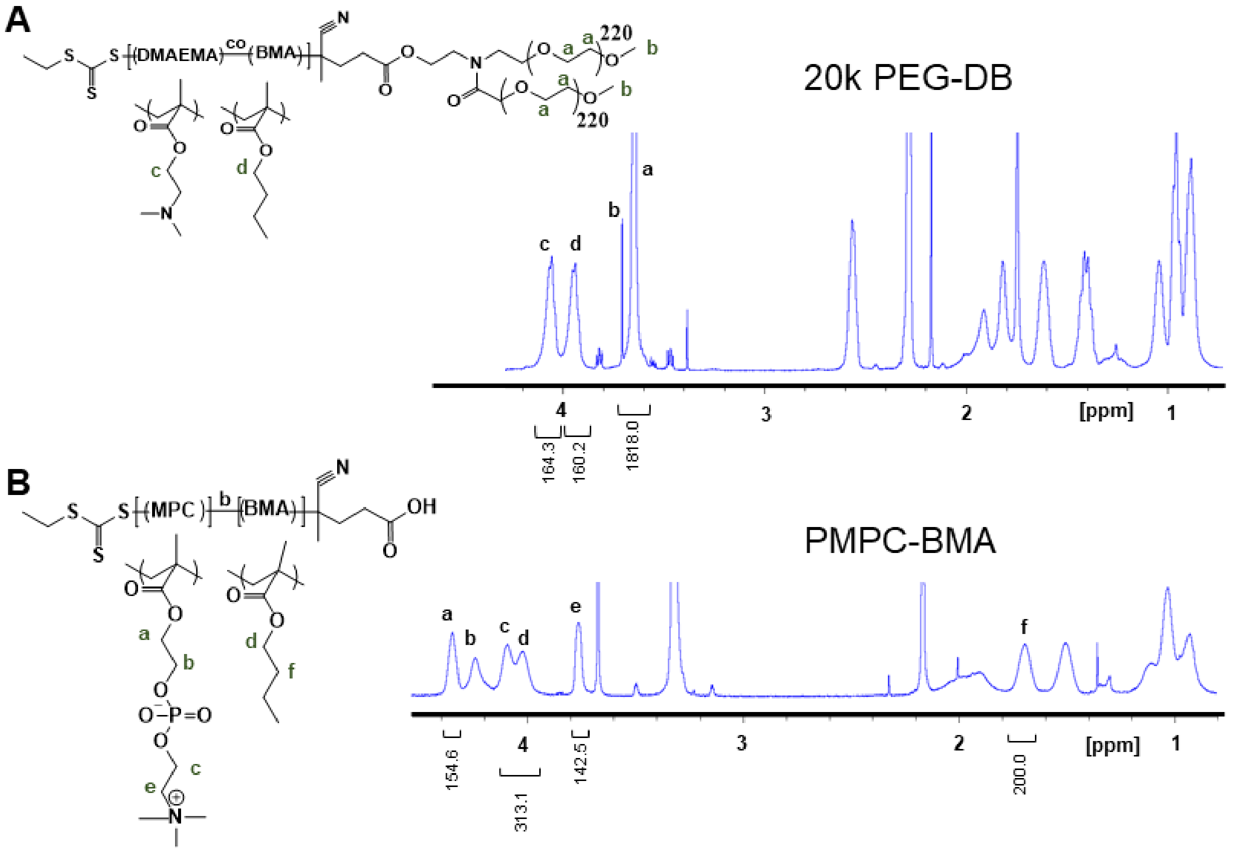


C

| Polymer | % DMAEMA:BMA in si-NP Core | Monomer DP | Total Mn (g/mol) |
|---|---|---|---|
| PMPC-DB (Zwitter si-NP) | 50:50 | MPC:70<br>D:78<br>B:78 | 42961 |
| PMPC-BMA (Zwitter Micelle) | 0:100 | MPC: 70<br>D:0<br>B:100 | 35149 |
| 20k PEG-DB (PEG si-NP) | 50:50 | D:80<br>B:80 | 43592 |

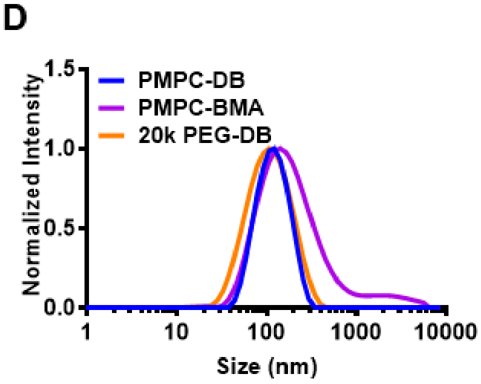


**Supplemental Figure S5**: si-NP characterization. [1]H-NMR characterizations for PEG si-NP, solvent MeOD (A) and zwitter micelle, solvent CDCl3 (B). C) Table of polymer characteristics, including Mn and DP. D) DLS curves of each si-NP formulation.

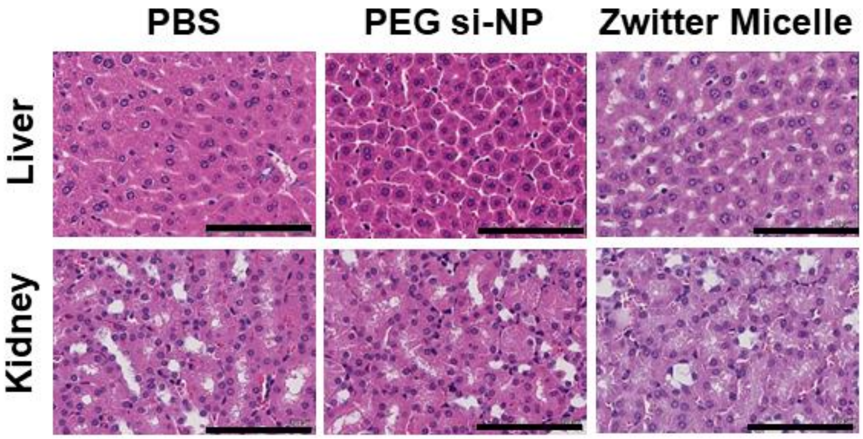


**Supplemental Figure S6**: Additional H&E staining for BALB/c mice treated with each si-NP formulation. Scale bars = 100 µm.

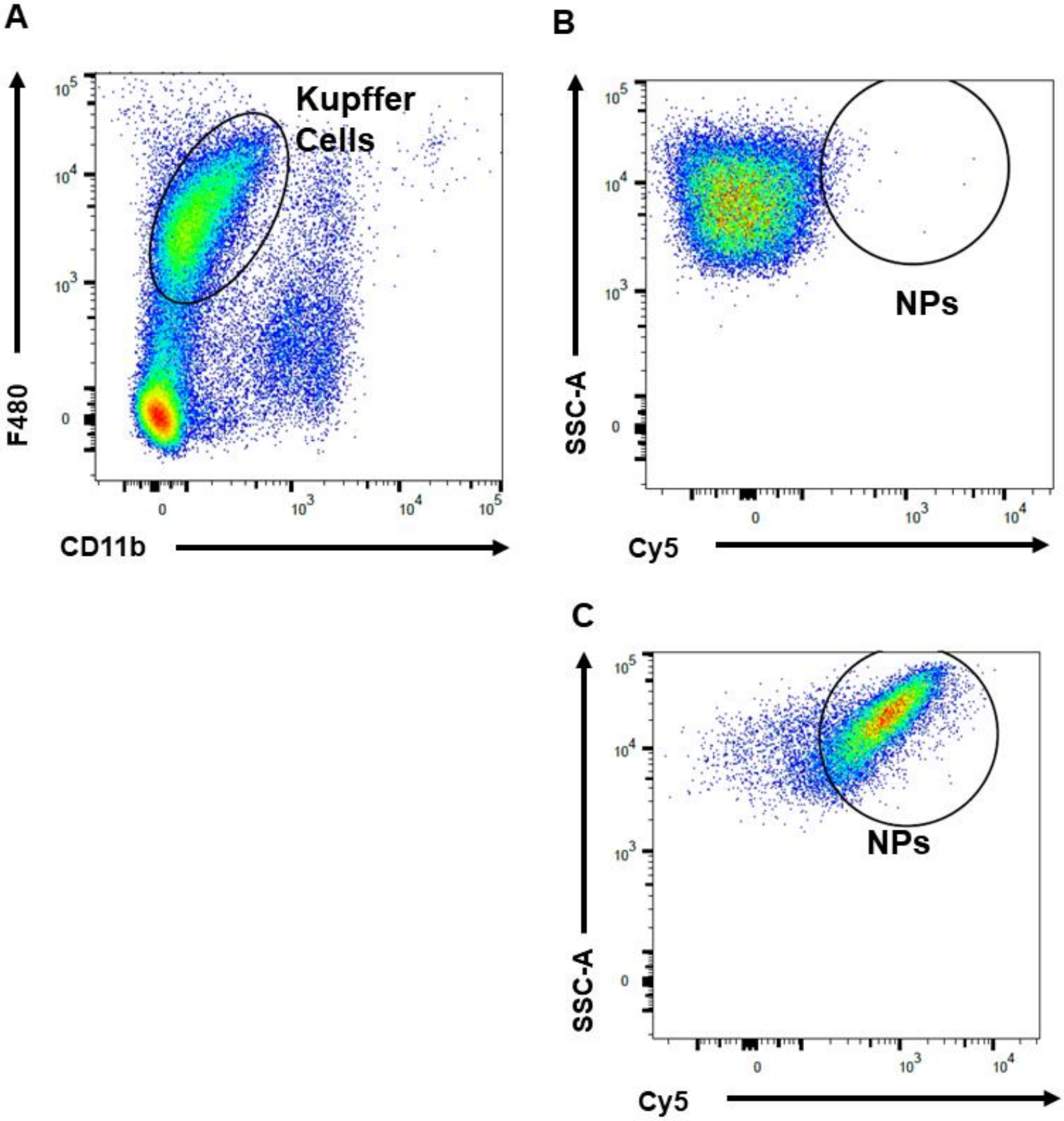


**Supplemental Figure S7.** Flow cytometry gating for measuring uptake of Cy5-labeled si-NPs by liver Kupffer cells of injected mice. After gating for live (DAPI-), singlet lymphocytes (CD45+), Kupffer cells were identified as F4/80+, CD11b low-intermediate (A). Kupffer cells positive for si-NPs were identified by Cy5+. Examples negative (B) and positive (C) for si-NP uptake are shown.

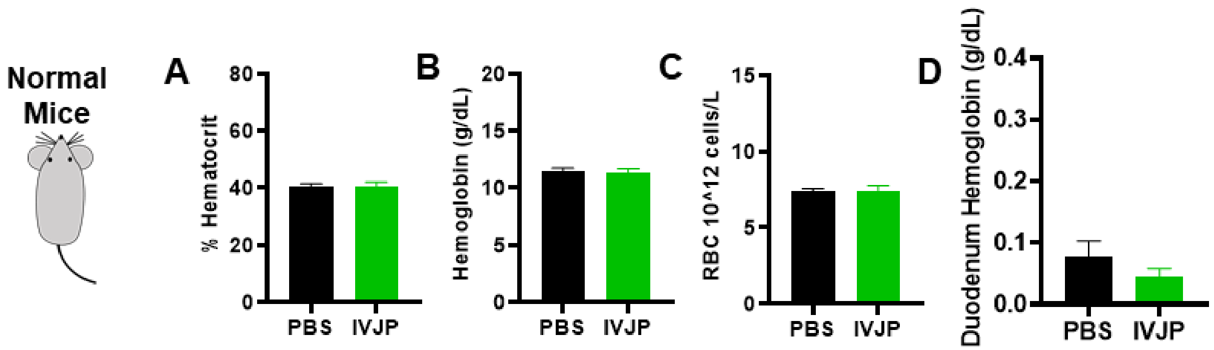


**Supplemental Figure S8**: Blood hematocrit (A), hemoglobin (B), red blood cell concentration (C), and duodenum hemoglobin concentration (D) for normal BALB/c mice treated with saline or 2 mg/kg IVJP.

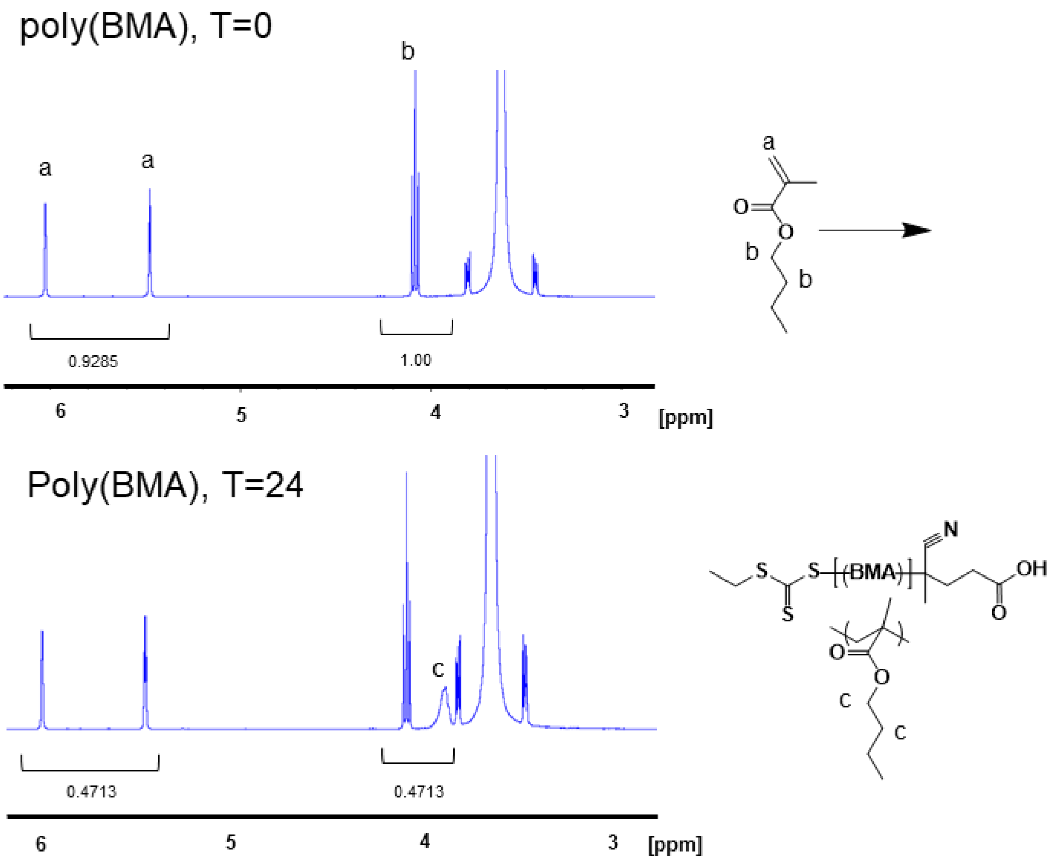


**Supplemental Figure S9**: $^{1}$H-NMR demonstrating synthesis of poly(BMA). Monomer conversion was calculated based on the disappearance of monomer peaks, as shown. $CDCl_3$ solvent.
